# Learning spatial cellular state through 3D reconstruction of the cell

**DOI:** 10.64898/2026.11.09.765489

**Authors:** Young Je Lee, Kanupriya Anad, Seonghui Min, Jose Lugo-Martinez

## Abstract

Subcellular-resolution spatial transcriptomics localizes individual RNA transcripts within cells with high-precision. Existing methods for identifying spatially organized cell populations rely on cellular niches defined by neighboring cells, leaving open whether such populations can be identified from individual-cell representations alone. Moreover, representations that jointly capture transcript identity and three-dimensional (3D) spatial organization have not been systematically learned or biologically evaluated. Here we present a representation learning framework that models each cell as a gene-annotated point cloud of transcripts and trains a rotation-equivariant neural network to reconstruct 3D cell shape, yielding spatially informed transcript embeddings. We further aggregate these embeddings into a cell-state representation that summarizes transcript-level spatial patterns. Our framework outperforms existing point-cloud-based 3D reconstruction methods on cell-shape reconstruction. In MERFISH human uterine cancer, combining the learned embeddings with gene expression improved prediction of CD45RO protein intensity over both gene expression alone and niche-based embeddings. Applied to MERFISH human liver cancer and Xenium human lung datasets, our framework identified distinct spatial patterns between cell populations, including spatially organized subpopulations marked by disease-associated gene expression and patterns associated with pathological tissue structures. Together, these results show that learning transcript representations through 3D cell reconstruction provides a novel approach to capture cell-state heterogeneity and biologically relevant spatial organization in subcellular spatial transcriptomics.

## 1 Introduction

The development of spatially resolved transcriptomics (SRT) has enabled gene expression to be examined within its tissue context. Among these technologies, high-resolution spatial transcriptomics approaches, such as MERFISH [1] and Xenium [2], localize individual transcripts in three-dimensional space with high precision. Spatial patterns in these data are commonly investigated by visualizing the spatial distributions of selected genes or by analyzing local cellular neighborhoods [3, 4]. One common strategy in niche analysis is to characterize the cellular neighborhood of each cell based on the composition of nearby cell types, followed by clustering to identify recurring spatial niches [5]. Building on this approach, several computational methods have been developed to generate cell embeddings that incorporate neighboring-cell information, enriching the analysis of spatial transcriptomics data [6, 7]. Although these approaches provide valuable insights into tissue organization, they primarily characterize spatial patterns using information from cellular neighborhoods. Because cellular phenotypes are influenced by their tissue microenvironment, molecular features of individual cells may also reflect aspects of their surrounding spatial context. We therefore asked whether spatially organized cell populations could be identified from molecular representations of individual cells without explicitly incorporating neighboring-cell information.

One opportunity to investigate this question lies in the spatial configuration of individual transcripts within cells. While conventional cell-by-gene expression profiles characterize transcript abundance, aggregating transcripts into gene counts removes their individual spatial coordinates, thus, potentially obscuring information about their organization within cells. High-resolution spatial transcriptomics provides both the gene identities and spatial positions of individual transcripts, offering an opportunity to construct cell-level representations using transcript-level information. Beyond spatial domain identification, evaluating learned representations through independent molecular prediction tasks provides a complementary assessment of their biological utility. Incorporating transcript-level spatial information and cellular shape into learned representations offers an additional opportunity to characterize individual cells. However, learning informative representations from these data presents several challenges. First, designing a meaningful learning objective is cumbersome because gene identity is already available as an input feature, making direct prediction of the same identity a potentially trivial task. Second, variation in cellular shape and orientation complicates the construction of representations that generalize across cells without relying on a fixed coordinate frame. Addressing these challenges requires an approach that learns spatial relationships among transcripts through an informative training objective while accounting for geometric variation across cells.

Representing transcripts as gene-labeled 3D point clouds provides a natural framework for addressing these challenges. Geometric equivariant neural networks can model spatial relationships among transcripts while accounting for transformations in cellular orientation. In computer vision, 3D reconstruction from point clouds has been extensively studied to learn geometric representations of objects [8, 9]. Building on this direction, 3D cell reconstruction from transcript point clouds provides a representation-learning objective that relates the spatial configuration of transcripts to cellular structure. By training a model to reconstruct 3D cellular structures, transcript embeddings can be learned through a geometric reconstruction objective rather than through direct prediction of gene identities. Furthermore, the reconstruction process provides an opportunity to learn representations associated with cellular geometry. Together, these learned representations can subsequently be used to characterize individual cells without explicitly incorporating neighboring-cell composition.

Here, we propose Spatial Cell State Selector (**SpaCState**), a computational framework designed to identify spatially organized cell populations using representations learned from individual cells. SpaCState employs an equivariant neural network to learn transcript embeddings through a 3D cell reconstruction objective, incorporating information about spatially neighboring transcripts within each cell. The framework also generates geometric embeddings through the reconstruction process. We further introduce *Spatialstate*, a continuous cell-level representation derived from learned transcript embeddings, to characterize individual cells and identify spatially organized cell populations. We benchmark the 3D reconstruction performance of SpaCState against existing point-cloud reconstruction methods. To independently evaluate the biological utility of the learned embeddings, we also assess their ability to predict CD45RO protein intensity in MERFISH human uterine cancer samples, demonstrating improved predictive performance compared with gene expression alone and embeddings from niche-based methods. We further apply SpaCState to MERFISH liver cancer and Xenium human lung datasets, identifying representations associated with distinct spatially organized cell populations, including macrophages surrounding necrotic regions and fibroblasts associated with fibrotic foci. Together, these results demonstrate the biological utility of reconstruction-derived cellular representations and the feasibility of identifying spatially organized cell populations using individual-cell representations without explicitly incorporating neighboring-cell information. st2sc is available on GitHub: https://github.com/youngje-cs-cmu/SpaCState

## 2 Method

In this section, we first define the problem, then introduce SpaCState, a computational framework that captures spatial patterns from individual cells using 3D reconstruction. Finally, we describe *Spatialstate* which captures the transcript-level spatial patterns into a cell state representation from high resolution spatial transcriptomics.

### 2.1 Problem formulation

We consider imaging-based spatial transcriptomics data, where each transcript is characterized by its gene identity and three-dimensional spatial coordinates. Let *N* denote the number of cells in a sample and *M* denote the number of distinct genes, indexed by *i* and *j*, respectively. Let *n*_*i j*_ denote the number of transcripts of gene *j* in cell *i*, and let *T*_*i jk*_ ∈ ℝ^3^ represent the spatial coordinates of the *k*-th transcript of gene *j* in cell *i*. Our objective is to learn a function *f*_*θ*_ that maps each transcript to a *D*-dimensional embedding vector,

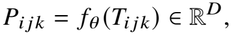

where *D* denotes the dimension of the transcript embedding. The learned representation *P*_*i jk*_ aims to capture both the gene identity of the transcript and the local spatial organization of neighboring transcripts. We subsequently use these transcript-level representations to characterize gene- and cell-level spatial organization.

### 2.2 Overview of SpaCState framework

Here, we propose SpaCState (Spatial Cell State Selector), a framework that learns spatial cellular states through 3D reconstruction. Using an equivariant neural network to reconstruct 3D cellular voxel from the 3D transcripts as a biological objective, the framework generates transcript embeddings that encode neighboring transcripts information. The input 3D transcripts first pass the transcript encoder and produce transcript embeddings (Fig. 1B). The transcript embeddings produced are then passed into the grid decoder to reconstruct the 3D cell voxel and produce grid embeddings. For interpretability, the transcript embeddings are then processed into *Genestate* and *Spatialstate*, described in Section 2.4, to capture the cellular state from spatial neighborhood information (Fig. 1C).

**Fig. 1.**
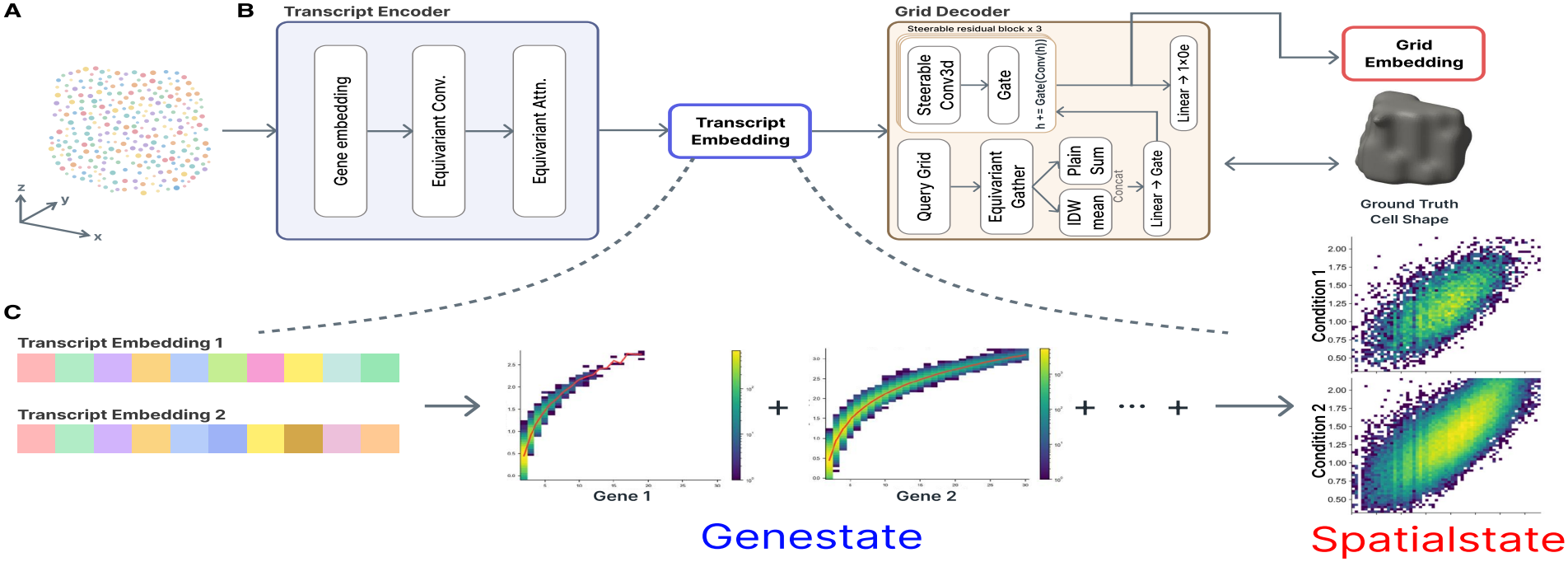
Overview of the SpaCState framework and Spatialstate construction. (A) Individual transcripts are represented as 3D point clouds, with spatial coordinates and gene identities. (B) The transcript encoder uses gene embeddings, equivariant convolution, and equivariant attention to generate transcript embeddings. These embeddings are subsequently passed to the grid decoder to reconstruct the 3D cellular structure, while also generating grid embeddings. (C) The learned transcript embeddings are used to construct gene-specific distributions, which are subsequently aggregated to generate cell-level representations under different conditions.

### 2.3 Equivariant neural network

To learn transcript representations that incorporate local spatial information, we employ a rotation-equivariant neural network based on the e3nn framework. We first encode the gene identity of each transcript as a type-*l* = 0 feature, denoted by 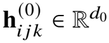, where *d*_0_ denotes the dimension of the gene identity encoding. Since gene identity is independent of spatial orientation, the type-*l* = 0 feature remains invariant under any rotation *R* ∈ *SO* (3), satisfying

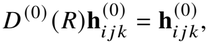

where *D* ^(0)^ *R* = 1 denotes the trivial representation of the rotation group, acting independently on each feature channel.

To incorporate spatial information, we construct a local neighborhood *N*_*K*_ (*T*_*i jk*_) consisting of the *K* nearest transcripts within the same cell, based on Euclidean distance. For each neighboring transcript 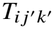, we compute the relative displacement vector

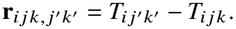

The equivariant neural network aggregates neighboring gene identity features and their relative spatial information through equivariant operations, allowing the learned transcript representation to incorporate local transcript organization while preserving rotational equivariance.

The resulting transcript embedding *P*_*i jk*_ is composed of type-*l* features, which we denote by 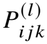. Let ℒ denote the set of irreducible representation orders used in the output embedding. We express the transcript embedding as

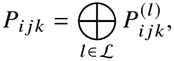

where ⨁ denotes concatenation of the type-*l* feature components. For each type *l*, the feature 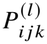 transforms according to its corresponding irreducible representation under spatial rotations.

### 2.4 Genestate and Spatialstate

The transcript embedding *P*_*i jk*_ incorporates both the gene identity of a transcript and information from its neighboring transcripts. Since transcripts with the same gene identity share the same initial type-*l* = 0 feature, we hypothesize that differences between their learned embeddings can reflect variations in their local spatial environments. Specifically, comparing transcripts of the same gene within a cell allows us to characterize differences in their neighboring transcript organization while holding gene identity fixed. Based on this hypothesis, we define *Genestate* to quantify within-gene variation in transcript embeddings.

For gene *j* in cell *i*, we consider all unique pairs of its *n*_*i j*_ transcripts and compute the Euclidean distance between their embeddings. We define the *Genestate* score *g*_*i j*_ as

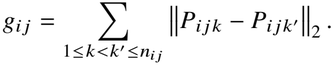

Since each transcript embedding consists of type-*l* features, the *Genestate* score can equivalently be expressed as

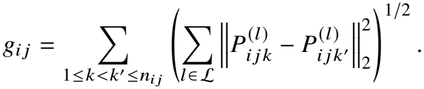

For genes with fewer than two transcripts in a cell, we define *g*_*i j*_ = 0.

Although the transcript embeddings may contain higher-order equivariant features, the Euclidean distances between corresponding embeddings are invariant under spatial rotations. Consequently, the *Genestate* score provides a rotation-invariant measure of within-gene transcript embedding variation.

We further define the *Spatialstate* vector of cell *i* by aggregating the *Genestate* scores across all *M* genes:

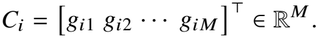

Thus, each element of *C*_*i*_ represents the aggregated pairwise embedding differences among transcripts of a particular gene. The resulting *Spatialstate* vector summarizes gene-specific variations in local transcript organization, providing a cell-level representation derived from the spatial relationships among individual transcripts.

To obtain a single cell-level score, we average the *Spatialstate* vector over the genes that have at least two transcripts in the cell. Let *M*_*i*_ denote the number of such genes in cell *i*. We define the *Spatialstate* score *c*_*i*_ as

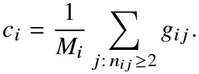

Pairwise embedding differences are therefore accumulated within each gene, so that a gene with more transcripts contributes a larger score, and then averaged across the genes present in the cell, so that *c*_*i*_ does not grow with the number of expressed genes alone. Genes with fewer than two transcripts, for which *g*_*i j*_ = 0, are excluded from the average.

The construction of the *Spatialstate* can be subjected by condition, such as cell type or disease state. The approach provides two ways to interpret spatial pattern in the spatial transcriptomics using *Spatialstate*. First way, use of *Spatialstate* itself. Display the *Spatialstate* spatially shows cells are spatially clustered based on the *Spatialstate*. Another way for interpretation is we construct 2D histogram from *Spatialstate*, where x axis monitors number of the transcripts, and y axis to view the distribution of each state value. This histogram shows how cells are distributed under the *Spatialstate* across varying number of the transcripts and summation of *Spatialstate*, which reflects the distribution of population of cells of same *Spatialstate*.

### 2.5 Reconstruction of the 3D cell from 3D transcripts and grid embedding

To learn spatially informative transcript embeddings for the construction of *Genestate* and *Spatialstate*, we employ 3D cell reconstruction as an representation-learning objective. Specifically, we train the equivariant neural network to extract transcript embeddings that support the reconstruction of three-dimensional cellular morphology from transcript point clouds. The reconstruction task serves as a supervisory objective for learning transcript representations, rather than as the primary goal of the proposed framework.

For each cell *i*, let *P*_*i*_ denote the set of transcript embeddings obtained from the equivariant neural network *f*_*θ*_. To reconstruct the three-dimensional cellular volume, we employ a grid-based reconstruction decoder that aggregates transcript embeddings onto a regular voxel grid.

Let **q***_V_*∈ℝ^3^ denote the spatial coordinates of voxel *v*, and let _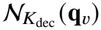_ denote the set of its *K*_dec_ nearest transcripts. For each voxel, we aggregate neighboring transcript embeddings and their relative spatial information to construct a voxel-level feature representation:

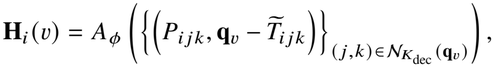

where *A*_*ϕ*_ denotes the equivariant spatial aggregation function and 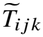 represents the normalized transcript coordinates. The resulting voxel-level features are arranged into a three-dimensional feature grid **H**_*i*_, which is processed through steerable 3D convolutional layers *s*_*ϕ*_ to incorporate spatial information across neighboring voxels, yielding an equivariant feature field on the grid,

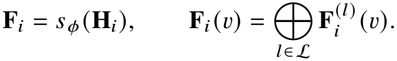

The predicted occupancy field is obtained by a linear read-out *w*_*ϕ*_ of the type-*l* = 0 component at each voxel,

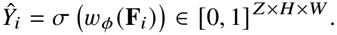

The grid embedding of cell *i* is obtained from **F**_*i*_ by reducing the voxel-level features to rotation invariant representations. Specifically, we retain the type-0 components as they are and the norm of every higher-order component, and averaging over the voxels of the segmented cell volume,

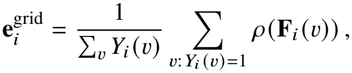

where *ρ* denotes the invariant reduction.

The reconstruction target is defined by a binary occupancy field

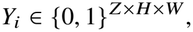

obtained from the segmented cellular volume. To train the transcript encoder and reconstruction decoder jointly, we minimize a combination of binary cross-entropy (BCE) and Dice losses:

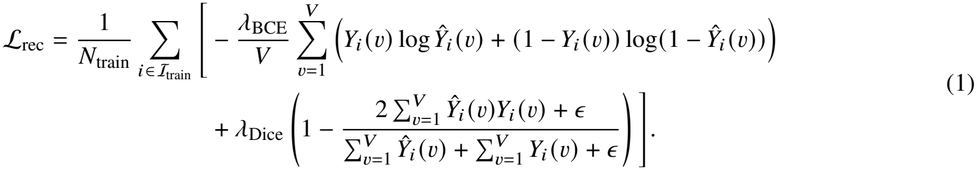

where *λ*_BCE_ and *λ*_Dice_ denote the weighting coefficients of the two loss terms.

For reconstruction evaluation, the predicted occupancy field is binarized at a threshold *τ*, and voxel-level intersection-over-union (IoU) is computed:

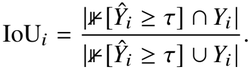

The learned transcript embeddings are subsequently used to construct *Genestate* and *Spatialstate* representations, independently of the reconstruction decoder.

### 2.6 Dataset preparation

Imaging-based spatial transcriptomics datasets provide transcript-level gene identities and three-dimensional spatial coordinates, together with microscopy images that can be used to obtain cellular segmentation masks. To construct the training data for the reconstruction objective, we first obtain three-dimensional cell segmentation masks from the corresponding microscopy images using Cellpose3 [10]. Each segmented region is converted into a binary occupancy field *Y*_*i*_.

To associate transcripts with their corresponding segmented regions, we align the transcript coordinates with the microscopy image coordinate system using a dataset-specific affine transformation. Following coordinate alignment, transcripts are associated with their corresponding segmented regions based on spatial overlap. For each cell *i*, we center and normalize the transcript coordinates. The corresponding segmentation mask is transformed into the same normalized coordinate system and represented on a regular voxel grid to obtain *Y*_*i*_. This ensures spatial consistency between the transcript point cloud and its reconstruction target.

For evaluation of the reconstruction objective, we used MERFISH human liver cancer from Vizgen MERFISH FFPE Human Immuno-oncology Data Set released in May 2022 and Xenium human breast cancer datasets[2]. For evaluation of the embeddings, we used MERSCOPE FFPE Anti-body Co-staining Data Release. For downstream *Genestate* and *Spatialstate* analyses, we used MERFISH human liver cancer and Xenium human lung datasets. The MERFISH datasets provided cell membrane staining images, while the Xenium datasets provided nuclear staining images. Three-dimensional segmentation masks were obtained from the corresponding images using Cellpose3. To evaluate with existing reconstruction methods, we used anisotropy option in Cellpose3 to be handled to make data near isotropic as possible, with anisotropy values were selected based on the dataset.

### 2.7 Evaluation of 3D cell reconstruction

To benchmark 3D reconstruction, we compared our model with two methods that reconstruct 3D surfaces from point clouds, Convolutional Occupancy Networks (ConvONet) [8] and the Implicit Autoencoder (IAE) [9], using each cell’s transcript coordinates as the input point cloud. ConvONet encodes the input point cloud into structured feature maps that are processed by convolutional networks. Its decoder then predicts, for any query point in space, the probability that the point lies inside the cell, so the cell is represented as an occupancy field. IAE, originally developed for self-supervised representation learning of point clouds, encodes the input points into a latent representation from which the shape is reconstructed. Rather than predicting occupancy, its decoder predicts the unsigned distance from each query point to the cell surface, so the cell is represented as a distance field. In computer vision, such reconstructions are typically evaluated using the Chamfer distance between points sampled from the reconstructed and ground-truth surfaces. Because the ground truth for each cell is a voxel mask, we instead converted each reconstructed surface into a voxel grid and evaluated reconstruction accuracy using voxel-wise IoU. Because both methods predict a continuous field (occupancy for ConvONet and distance for IAE) rather than a voxel grid, we sampled the predicted field on a 643 grid, thresholded it to define the cell surface (at an occupancy probability of 0.2 for ConvONet and, for IAE, at the distance value that best separated interior from exterior voxels), mapped this surface into the ground-truth voxel frame, and computed the IoU between the voxels it enclosed and the ground-truth mask.

### 2.8 Evaluation of embeddings

To evaluate the embeddings learned through reconstruction, we used the MERSCOPE FFPE Antibody Co-staining Data Release. The sample consists of a human uterine cancer tissue section measured using MERFISH with a 500-gene panel, together with CD45RO, a marker of T-cell differentiation and activation. Both the cell membrane and CD45RO antibody images consist of seven slices, allowing for 3D cell reconstruction and 3D intensity measurements. In this evaluation, we included NicheScope [5], CellCharter [7], and BANKSY [6], which are niche identification methods for spatial transcriptomics that generate cell embeddings. For each cell segmented using Cellpose3 and annotated as a lymphocyte, we computed the total CD45RO intensity within the cell mask in the intensity image, which shares the same coordinate space as the segmentation image. The lymphocytes were randomly split into training and testing sets: 75% (17,946 cells) were used to train a Ridge regression model using each embedding or gene-expression profile alone as an independent baseline to predict the log_10_-transformed total intensity, and 25% (5,983 cells) were used for testing. To assess the correlation between the embeddings and CD45RO intensity, we computed the Pearson correlation coefficient between the predicted and measured intensities of the test cells. To evaluate CD45RO prediction performance, we calculated *R*^2^ on the test cell

### 2.9 Cell type annotation

Cell type annotation for the datasets used in downstream analyses was performed using Ingest from Scanpy [11]. For the evaluation of the embeddings from SpaCState, Regner et al. [12]. were used to annotate cells from human uterine cancer sample. For the MERFISH liver cancer data analysis, Sharma et al. [13] and Ma et al. [14] were used. For the Xenium human lung analysis, the Human Lung Cell Atlas was used [15]. Due to the size of the reference dataset, we first filtered out cell types with fewer than 200 cells and those that were difficult to interpret (Submucosal gland, None, and Rare), retaining 18 cell types at level 3, and then subsampled the dataset to 150,000 cells.

### 2.10 Application on MERFISH human liver cancer

The MERFISH liver cancer samples used in the reconstruction evaluation were also used in this analysis. Two replicates measured with the same 500-gene panel were used in this study. The enrichment of T cell activation (GO:0042110) was computed using GSEApy [16]. To identify differentially expressed genes between macrophages with a high cell state and the other macrophages, a log10 cell-state threshold of 1.68 was applied, and a two-sided Mann-Whitney U test was performed on the log-normalized expression of each gene.

### 2.11 Application on Xenium human IPF lung

In this study, we used the Xenium dataset of Vannan et al. [17], which comprises 16 idiopathic pulmonary fibrosis (IPF) samples and 10 controls. The dataset includes pathologist annotations of cells within fibrotic foci. We also connected these annotated cells to define the region boundaries. We tested two hypotheses: (i) fibroblasts in IPF samples have higher *Spatialstate* inside annotated fibrotic foci than outside them, and (ii) SPP1+ and FABP4+ macrophages have higher *Spatialstate* than other macrophages. For each hypothesis, we performed a one-sided sign test separately within each sample.

## 3 Results

In this section, we show that our SpaCState is competent against existing point cloud-based object reconstruction, and shape-encoded embeddings can benefit downstream biological analyzes through evaluation on MERFISH human uterine cancer samples. Next, we demonstrate that SpaCState can detect transcript-splitting problems and capture spatial patterns of macrophages in MERFISH liver cancer samples, as well as fibroblasts and disease-associated macrophages in Xenium human lung samples.

### 3.1 SpaCState reconstructs 3D cellular shape from 3D transcript point clouds

We first evaluated our 3D reconstruction performance. Since our learning dependency on the reconstruction of the 3D cell as biological objective, naturally, reconstruction performance can be assessed. In particular, we evaluated our method against 3D point cloud reconstruction models for cell and nucleus reconstruction tasks (See Methods for details). In both cell reconstruction from MERFISH data and nucleus reconstruction from Xenium data, our method showed favorable performance against established object reconstruction methods (Supp. Table I).

### 3.2 Shape-encoded embeddings from SpaCState predict CD45RO expression in MERFISH uterine cancer

Next, We evaluated whether the embeddings learned by SpaCState capture biologically meaningful representations through the reconstruction task. To maximize the benefits of the learned embeddings, we used both transcript embeddings and grid embeddings from the encoder and decoder, respectively. The MERSCOPE FFPE Antibody Co-staining Data Release program includes CD45RO co-staining data from human uterine cancer samples (Fig. 2A). CD45RO measurements provide meaningful information about where T-cell activity is spatially enriched, allowing us to better understand T-cell activity in cancer. This led us to investigate whether the learned embeddings could predict CD45RO intensity in T cells. Because the dataset provides lymphocyte annotations at a fine-grained level, we evaluated the correlation between the learned embeddings and CD45RO intensity in lymphocytes as the first task, and the *R*^2^ of CD45RO intensity predictions from the learned embeddings as the second task (Fig. 2B-D). In both tasks, models trained using SpaCState embeddings outperformed existing methods, demonstrating that our embeddings captured biological information. Moreover, comparisons with gene expression showed that geometric information could further improve biological prediction performance compared with approaches using gene expression alone.

**Fig. 2.**
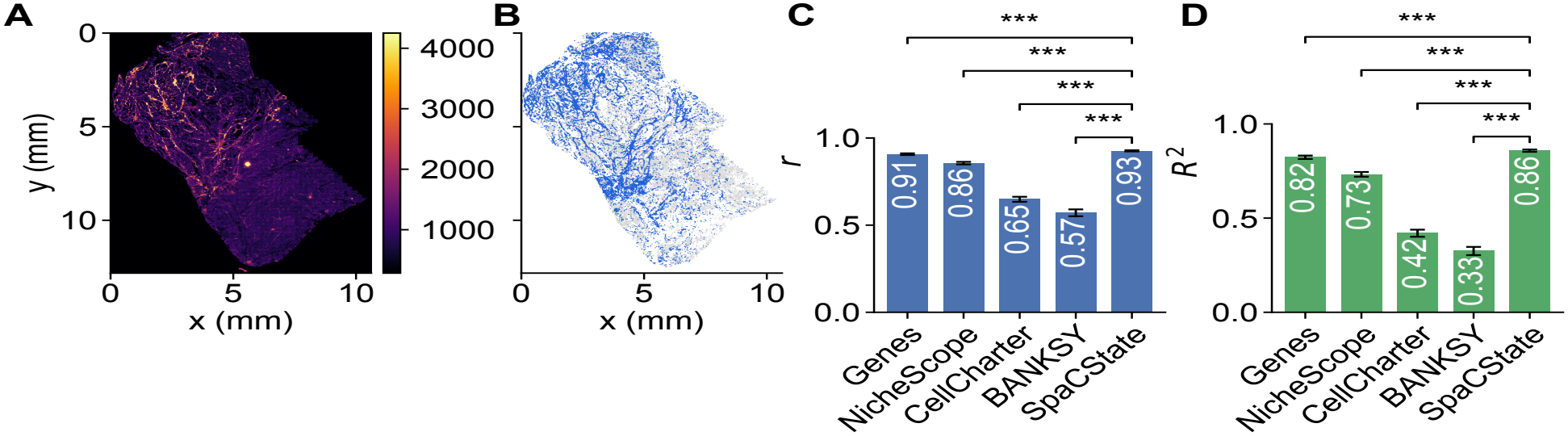
Shape-encoded embeddings improve the prediction of CD45RO intensity. (A) Intensity map of CD45RO. (B) Cell-type annotations, where blue cells represent lymphocytes and gray cells represent other cell types. (C) Bar plot showing the correlation between embeddings and CD45RO intensity using Pearson’s correlation coefficient. (D) Bar plot showing the prediction performance of CD45RO intensity using Coefficient of determination (*R*^2^).

### 3.3 SpaCState detects transcript splitting and biological domains of macrophages in MERFISH liver cancer

Next, to examine how our *Spatialstate* reflects spatial cellular states, we applied our framework to MERFISH human liver cancer samples. For cell-type annotation, we used two reference single-cell RNA-seq datasets due to their limited cell type coverage. Specifically, Sharma et al [13]. reference includes hepatocytes but lacks malignant cells, whereas Ma et al [14]. reference includes malignant cells but lacks hepatocytes. When we examined T cells, we observed different *Spatialstate* histograms were observed between two references. While the Sharma et al. reference showed two distinct highlighted regions, the Ma et al. reference showed only one highlighted region (Fig. 3A,B). We further investigated these two regions and observed that T cells in state 1 annotated using the Sharma et al. reference were annotated as malignant cells using the Ma et al. reference, whereas T cells in state 2 were reannotated as T cells (Fig. 3B). T cell activity enrichment analysis further showed significantly lower enrichment in state 1 than in state 2 (Fig. 3D). We hypothesize that this difference arises from transcript splitting problem in subcellular spatial transcriptomics, where inaccurate cell segmentation can lead to misleading biological interpretations. These results demonstrate that *Spatialstate* can distinguish cell populations with potentially inaccurate transcript assignments, highlighting its utility in detecting transcript-splitting artifacts.

**Fig. 3.**
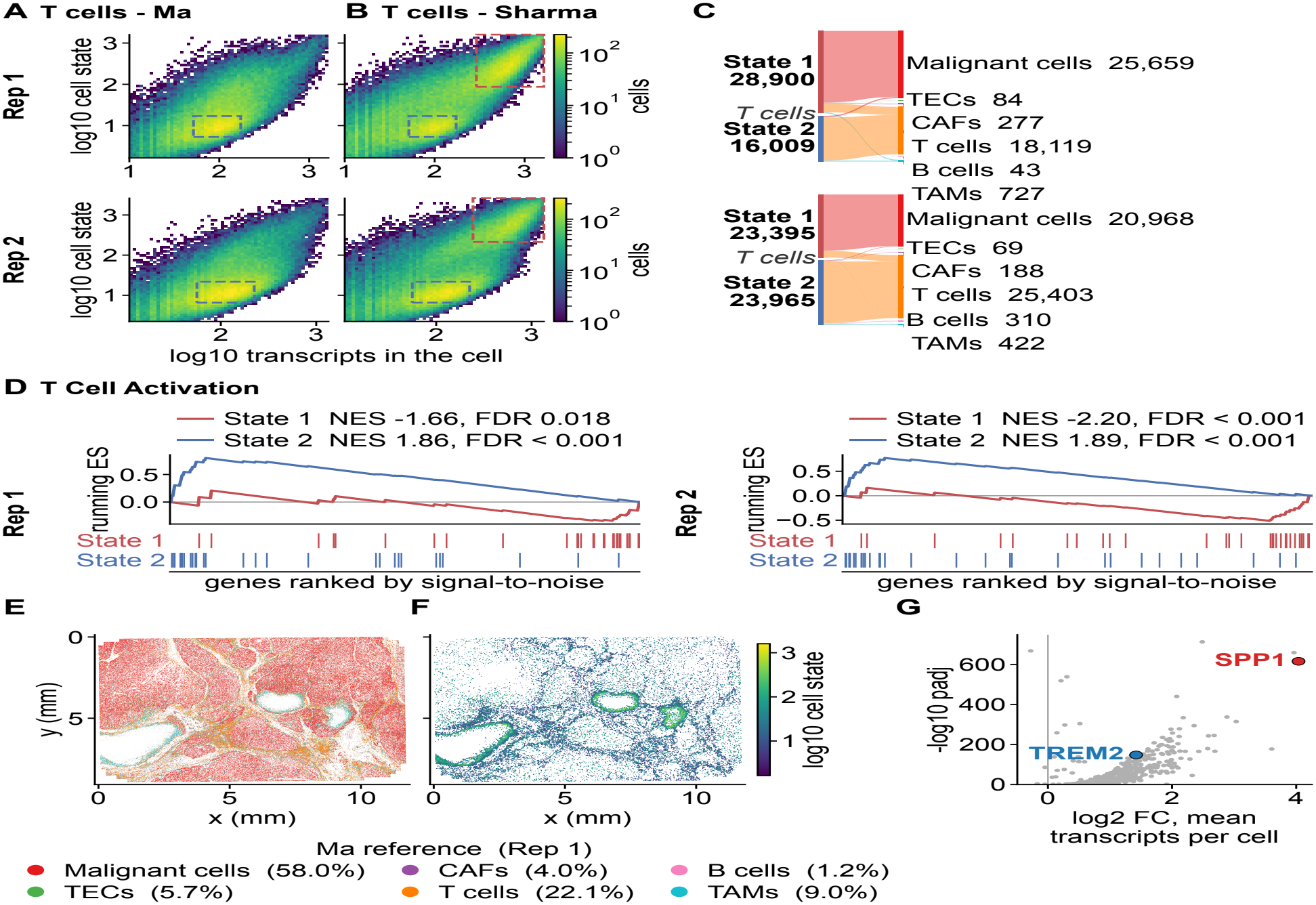
SpaCState detects transcript splits and a biological domain of macrophages in MERFISH liver cancer. (A) 2D histogram of the *Spatialstate* of T cells from the Ma reference. (B) 2D histogram of the *Spatialstate* of T cells from the Sharma reference. Red and blue boxes mark State 1 and State 2, defined on the Sharma T cells; the same boxes are shown in (A). (C) Sankey diagram showing the cell-type shift of T cells in the two states from the Sharma reference to the Ma reference. (D) Enrichment of T cell activation between T cells in the two states. (E) Cell-type annotation of replicate 1 using the Ma reference. (F) Spatial dot plot showing the *Spatialstate* of macrophages. (G) Volcano plot showing differentially expressed genes between the selected high-*Spatialstate* macrophages and the other macrophages.

Based on the comparison described above, we selected the Ma et al. reference for subsequent analysis. After cell type annotation, we observed that macrophages formed spatial clusters resembling rims around three distinct regions (Fig. 3E). Macrophages are known to form spatial rims around tumor boundaries [18], but these regions appeared to have few cells in their interiors. When we visualized the *Spatialstate* values of macrophages, we observed three regions forming spatial rims with higher *Spatialstate* values (Fig. 3F). The spatial organization of these regions is consistent with findings from Sopa [19], where these spatial clusters were detected by examining neighboring cells in necrotic domains. Necrotic regions initiate TREM2+ macrophages [20], and we were able to confirm high TREM2 expression in macrophages around the spatial rims, as reported by both Wang et al. and Sopa. However, while previous studies highlighted TREM2 expression in these regions, differential expression analysis of macrophages identified through our *Spatialstate* revealed elevated SPP1 expression without prior selection of marker genes (Fig. 3G). Given the heterogeniety of SPP1+ and TREM2+ macrophage populations, differential expression analysis of spatial clusters identified through our *Spatialstate* provides a complementary perspective on macrophage heterogeneity. This demonstrates that our framework captures these distinct biological domains through high *Spatialstate* values without examining neighboring cells.

### 3.4 SpaCState captures fibrotic foci and spatial clusters of disease-associated macrophages in Xenium IPF lung samples

Idiopathic pulmonary fibrosis (IPF) is a progressive lung disease characterized by fibrosis of unknown cause. As the disease progresses, honeycomb-like fibrotic structures develop, and the resulting scar tissue reduces respiratory function. Furthermore, SPP1+ macrophages and FABP4+ macrophages form spatial clusters in fibrotic lung samples [17]. The accumulation of SPP1+ macrophages is detected in the airways and increases as the disease progresses, whereas the extent of accumulation is less pronounced for FABP4+ macrophages.

We applied SpaCState to human Xenium IPF lung samples to test whether SpaCState could identify these spatial clusters using nucleus-level data. We used author-provided, expert-annotated fibrotic regions, including fibroblastic foci and severe fibrosis. (Fig. 4A). After training our model to reconstruct nuclei in 3D and extracting transcript embeddings, we constructed the *Spatialstate* representations of IPF fibroblasts. We first compared the 2D *Spatialstate* histograms of fibroblasts from healthy and IPF samples (Supp. Fig. 1A). Compared with healthy fibroblasts, IPF fibroblasts occupied a broader region of the 2D *Spatialstate* histogram. Notably, we observed increased *Spatialstate* histogram density in the upper-right region. When we examined disease-associated genes (CTHRC1, FAP, and POSTN), fibroblasts positive for these genes were also skewed toward the upper-right region. Based on these observations, we hypothesized that disease-associated fibroblasts exhibit higher *Spatialstate* values. When we displayed *Spatialstate* values in individual samples, we consistently observed that fibroblasts with high *Spatialstate* values were spatially asscoiated with fibrotic regions across multiple samples (Fig. 4B). A quantitative comparison of *Spatialstate* values between fibroblasts within fibrotic foci and other fibroblasts also supported this observation (Fig. 4C). These results suggest that *Spatialstate* can identify fibroblasts associated with fibrotic foci, highlighting regions of interest in IPF lung samples.

**Fig. 4.**
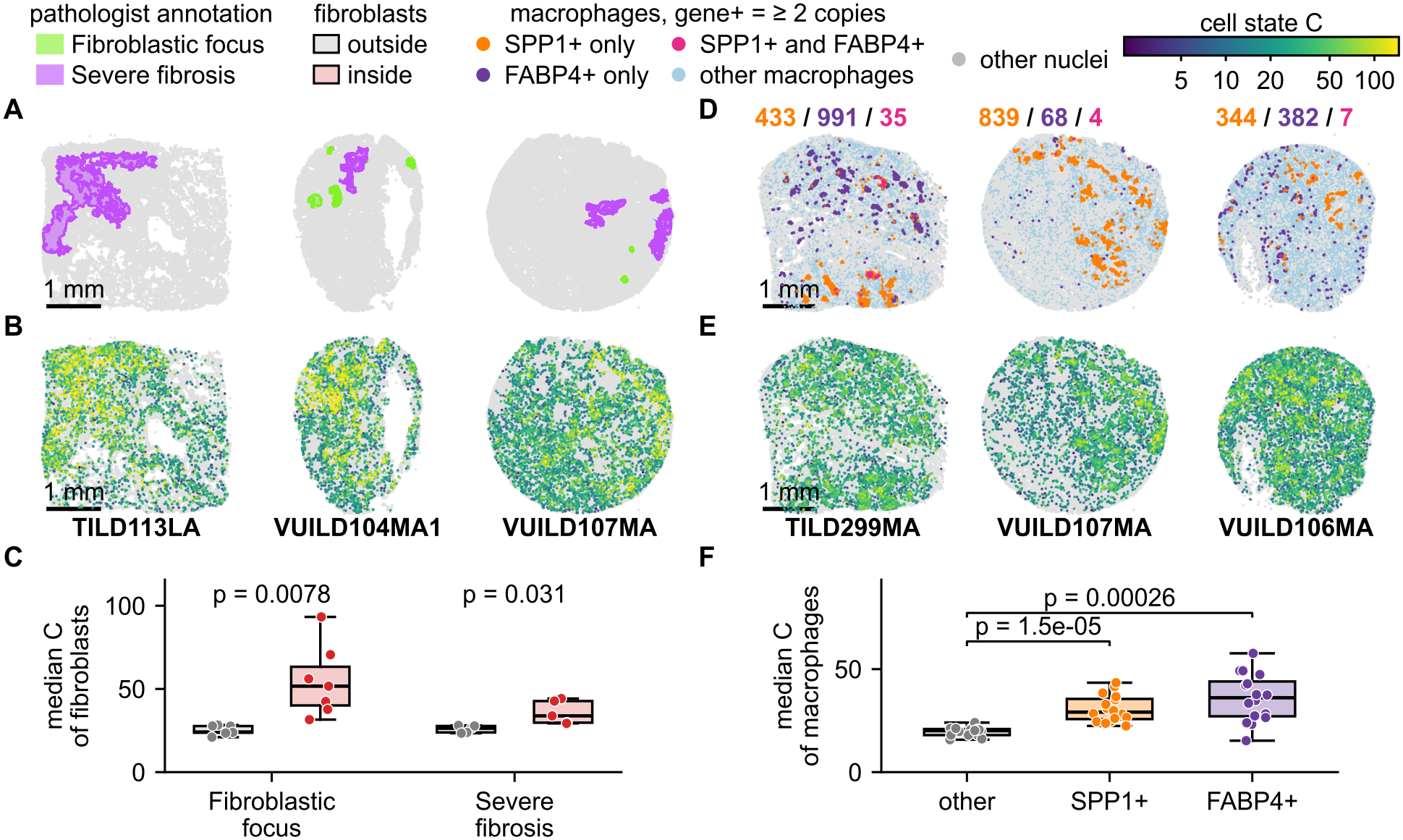
SpaCState captures fibrotic foci and spatial clusters of disease-associated macrophages in Xenium IPF lung samples. (A) Expert annotation of the fibrotic foci in IPF lung. (B) Spatial dot plot showing the *Spatialstate* values of fibroblasts. (C) Box plot showing the distribution of *Spatialstate* values for fibroblasts inside and outside the fibrotic foci. (D) Spatial dot plot showing the positions of disease-associated gene-positive macrophages. (E) Spatial dot plot showing the *Spatialstate* values of macrophages. (F) Box plot showing the distribution of *Spatialstate* values across SPP1+ macrophages, FABP4+ macrophages and other macrophages.

We then extended our analysis to macrophages. Unlike fibroblasts, macrophages showed similar *Spatialstate* distributions between healthy and IPF samples (Supp. Fig. 1B). Macrophages expressing disease-associated genes showed a less pronounced shift toward the upper-right region of the 2D *Spatialstate* histogram. When we displayed the *Spatialstate* values of macrophages, we observed spatial clusters exhibiting different *Spatialstate* values (Fig. 4E). This suggests that spatial clusters associated with distinct cell states can be identified across different *Spatialstate* values, rather than being restricted to high values. We then examined the relationships between *Spatialstate* values and SPP1+ and FABP4+ macrophage populations (Fig. 4D). Unlike fibrotic foci, which were defined by expert annotations of tissue regions, SPP1+ and FABP4+ macrophages were identified within the macrophage population, allowing us to directly compare *Spatialstate* values across macrophage subtypes. Both SPP1+ and FABP4+ macrophages showed *Spatialstate* values that differed significantly from those of other macrophages (Fig. 4F). Together with their spatial distributions, these results suggest that *Spatialstate* can distinguish disease-associated macrophage populations exhibiting distinct spatial patterns.

We also observed that spatial patterns could also be identified through the *Spatialstate* histogram. We observed a pronounced spike in the *Spatialstate* histogram of IPF multiciliated cells from IPF samples (Supp. Fig. 1C). Mapping cells corresponding to this spike back to their original tissue locations revealed spatial clusters of multiciliated cells in four samples (Supp. Fig. 1D). This observation suggests that spatially organized cell populations can be identified using both *Spatialstate* values and distinct features of the corresponding histograms. Together, these results demonstrate that *Spatialstate* identifies spatially organized cell populations across different cell types in human IPF lung samples without explicitly incorporating neighboring-cell information.

## 4 Discussion

Cells and their microenvironments interact bidirectionally. Characterizing spatial organization through cellular neighborhoods raises the question of whether similar spatial patterns can also be identified from individual cells. In this study, we proposed SpaCState, a computational framework that characterizes spatial cellular states using the learned *Spatialstate* representation. Our evaluation of CD45RO protein-intensity prediction further demonstrated the biological utility of representations learned through 3D reconstruction. By combining transcript-derived and geometric embeddings with gene expression, SpaCState improved prediction performance compared with gene expression alone. These findings suggest that reconstruction-derived representations contain information relevant to molecular characteristics of individual cells, supporting their use in downstream biological analyses. We further demonstrated that spatial patterns previously identified through neighborhood-based analyses can also be captured from individual-cell representations from individual cells using our framework.

While we demonstrated that *Spatialstate* scores can identify spatially organized cell populations, it remains unclear how best to determine appropriate thresholds. Furthermore, analyses of SPP1+ and FABP4+ macrophages revealed that spatially organized populations can exhibit distinct ranges of *Spatialstate* values, suggesting that a single threshold may not be sufficient to identify biologically meaningful cell populations. This also applies to the 2D histograms of *Spatialstate*. Currently, these patterns are identified through visual inspection. Future computational approaches could enable automated detection of histogram patterns indicative of spatially organized cell populations approach to detecting patterns indicative of spatial clusters.

Another direction for improving the characterization of spatial organization is to find an interpretable way to describe features of type *l* > 0. Although our framework incorporates higher-order equivariant features into the learned embeddings, interpreting their individual contributions to *r*emains challenging. Developing interpretable representations of higher-order features (*l* > 0) may provide further insights into the spatial organization of transcripts and their contributions to cellular states.

## 5 Acknowledgement

This work was partially supported by National Institutes of Health (NIH) grant 1U54AG075931 and National Science Foundation (NSF) award 2134998 (to J.L.-M.).

## References

[1] Kok Hao Chen et al. “Spatially resolved, highly multiplexed RNA profiling in single cells”. In: Science 348.6233 (2015), aaa6090.

[2] Amanda Janesick et al. “High resolution mapping of the breast cancer tumor microenvironment using integrated single cell, spatial and in situ analysis of FFPE tissue”. In: Biorxiv (2022), pp. 2022–10.

[3] Ruben Dries et al. “Advances in spatial transcriptomic data analysis”. In: Genome research 31.10 (2021), pp. 1706–1718.

[4] Giovanni Palla et al. “Spatial components of molecular tissue biology”. In: Nature biotechnology 40.3 (2022), pp. 308–318.

[5] Xinyi Yu et al. “NicheScope: Identifying Multicellular Niches and Niche-Regulated Cell States in Spatial Transcriptomics”. In: bioRxiv (2025), pp. 2025–08.

[6] Vipul Singhal et al. “BANKSY unifies cell typing and tissue domain segmentation for scalable spatial omics data analysis”. In: Nature genetics 56.3 (2024), pp. 431–441.

[7] Marco Varrone et al. “CellCharter reveals spatial cell niches associated with tissue remodeling and cell plasticity”. In: Nature genetics 56.1 (2024), pp. 74–84.

[8] Songyou Peng et al. “Convolutional occupancy networks”. In: european conference on computer vision. Springer. 2020, pp. 523–540.

[9] Siming Yan et al. “Implicit Autoencoder for Point-Cloud Self-Supervised Representation Learning”. In: 2023 IEEE/CVF International Conference on Computer Vision (ICCV). 2023, pp. 14484–14496. doi: 10.1109/ICCV51070.2023.01336.

[10] Carsen Stringer and Marius Pachitariu. “Cellpose3: one-click image restoration for improved cellular segmentation”. In: Nature methods 22.3 (2025), pp. 592–599.

[11] F Alexander Wolf, Philipp Angerer, and Fabian J Theis. “SCANPY: large-scale single-cell gene expression data analysis”. In: Genome biology 19.1 (2018), p. 15.

[12] Matthew J Regner et al. “A multi-omic single-cell landscape of human gynecologic malignancies”. In: Molecular cell 81.23 (2021), pp. 4924–4941.

[13] Ankur Sharma et al. “Oncofetal reprogramming in tumour development and progression”. In: Nature Reviews Cancer 22.10 (2022), pp. 593–602.

[14] Lichun Ma et al. “Single-cell atlas of tumor cell evolution in response to therapy in hepatocellular carcinoma and intrahepatic cholangiocarcinoma”. In: Journal of hepatology 75.6 (2021), pp. 1397–1408.

[15] Lisa Sikkema et al. “An integrated cell atlas of the lung in health and disease”. In: Nature medicine 29.6 (2023), pp. 1563–1577.

[16] Zhuoqing Fang, Xinyuan Liu, and Gary Peltz. “GSEApy: a comprehensive package for performing gene set enrichment analysis in Python”. In: Bioinformatics 39.1 (2023), btac757.

[17] Annika Vannan et al. “Spatial transcriptomics identifies molecular niche dysregulation associated with distal lung remodeling in pulmonary fibrosis”. In: Nature genetics 57.3 (2025), pp. 647–658.

[18] Yao Liu et al. “Identification of a tumour immune barrier in the HCC microenvironment that determines the efficacy of immunotherapy”. In: Journal of hepatology 78.4 (2023), pp. 770–782.

[19] Quentin Blampey et al. “Sopa: a technology-invariant pipeline for analyses of image-based spatial omics”. In: Nature Communications 15.1 (2024), p. 4981.

[20] Peng-Xiang Wang et al. “Multi-omic profiling identifies TREM2+ lipid-laden macrophages as inflammatory drivers and therapeutic targets of colorectal cancer liver metastasis”. In: Cancer Letters 656 (2026), p. 218657.

