## Supplementary document for "Learning spatial cellular state through 3D reconstruction of the cell"

### Appendix

**Supp. Table 1.** Evaluation of 3D cell reconstruction. All cells from one replicate were used for training, while cells from another replicate were used for testing.

| | MERFISH Liver Cancer | | Xenium Breast Cancer | |
|---|---|---|---|---|
| | Rep1->Rep2 | Rep2->Rep1 | Rep1->Rep2 | Rep2->Rep1 |
| ConvoNet | 0.661 | 0.650 | 0.480 | 0.490 |
| IAE | 0.735 | 0.692 | 0.488 | 0.528 |
| **SpaCState** | **0.754** | **0.703** | **0.586** | **0.590** |

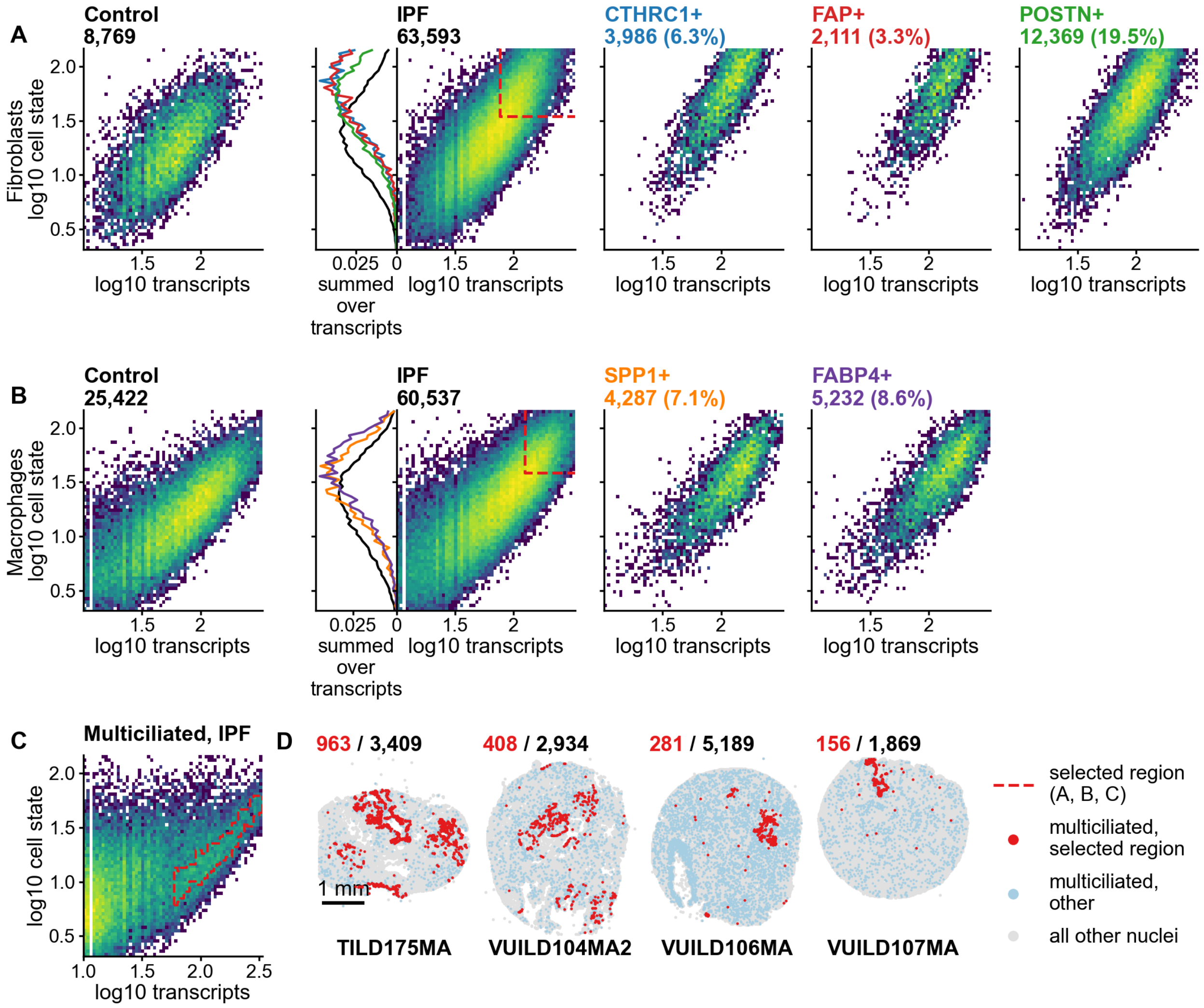


**Supp. Fig. 1. Extended figure of Xenium IPF lung analysis** (A) *Spatialstate* histograms of fibroblasts from control and IPF samples. (B) *Spatialstate* histograms of macrophages from control and IPF samples. (C) *Spatialstate* histograms of multiciliated cells from IPF samples. (D) Spatial dot plot showing the positions of multiciliated cells selected from red region in *Spatialstate* histogram.
